# Comprehensive evaluation of lifespan-extending molecules in *C. elegans*

**DOI:** 10.1101/2024.06.24.600458

**Authors:** Grace B. Phelps, Jonas Morin, Carla Pinto, Lucas Schoenfeldt, Sebastien Guilmot, Alejandro Ocampo, Kevin Perez

## Abstract

The nematode *C. elegans* has long served as a gold-standard model organism in aging research, particularly since the discovery of long-lived mutants in conserved aging pathways including daf-2 (IGF1) and age-1 (PI3K). Its short lifespan and small size make it highly suitable for high throughput experiments. While numerous molecules have been tested for their effects on *C. elegans* lifespan, consensus is still lacking regarding the most effective and reproducible compounds. Confounding effects, especially those related to drug-bacteria interactions, remain a contentious issue in the literature. In this study, we evaluated 16 of the most frequently reported lifespan-extending molecules in *C. elegans*, examining their effects on lifespan with two different diets (live and UV-killed OP50). In addition, we assessed the compounds’ impact on bacterial growth, their effects on various nematode strains, and the impact of the starting age of treatment. Our findings first confirmed robust lifespan extension with many, but not all, of the 16 tested compounds from the literature, and revealed that some of them could be combined to get synergistic effects. Additionally, we showed that some of these compounds also extend lifespan in the fly *D. melanogaster,* demonstrating a conserved effect across species. Finally, by expanding our screen to a broader pool of molecules, we identified novel lifespan-extending compounds in *C. elegans*.

## Introduction

The incidence of age-related diseases has dramatically risen due to an increasingly aging population, posing a significant burden on society and health care systems. It is therefore essential to identify therapeutic strategies that can prevent the functional decline associated with aging, the occurrence of age-related diseases and delay mortality in the elderly. The nematode *C. elegans* has a long history of being a gold standard model organism for aging research, notably with the identification of long-lived mutants in conserved aging pathways including daf-2^[1]^ (IGF1) and age-1^[2]^ (PI3K). Its short lifespan and small size make it amenable for large scale experiments. Although many compounds have been tested for their effect on *C. elegans* lifespan, there is still a lack of consensus on which ones are the best and most reproducible^[3,4]^. Confounding effects are also largely debated in the literature, notably concerning the interaction between the compound and the bacteria that are being fed to the worms. Recent efforts like the *C. elegans* intervention testing program (CITP^[5]^), counterpart of the mouse intervention testing program (ITP^[6]^), have attempted to standardize testing of lifespan-extending compounds in *C. elegans*, but so far with a relatively low number of compounds tested. Other approaches like the WormBot^[7]^ may facilitate higher throughput experiments in worms.

In this work, we surveyed 16 of the most extensively reported lifespan-extending compounds in *C. elegans*. We tested their effect on lifespan using two different types of food (live and UV-killed *E. coli* OP50). Moreover, we assessed the effect of the compounds on the bacteria itself, as well as their effect on lifespan of different nematode strains, and at multiple ages. In addition, we identified several new combinations of compounds with synergistic effect on lifespan. Importantly, we also showed that some of these compounds extend lifespan in *D. melanogaster,* demonstrating a conserved effect across species. Lastly, we opened our screen to a larger pool of compounds and identified novel lifespan-extending molecules in *C. elegans*.

## Results

### Effect on lifespan of 16 of the most frequently reported lifespan-extending compounds in C. elegans fed live OP50

We first sought to validate multiple lifespan-extending compounds reported in the literature. For this we surveyed DrugAge^[8]^, a database for lifespan extension data in different model organisms, and further performed a manual search. This yielded an extensive list of putative lifespan-extending compounds in the *C. elegans* strain N2. We specifically focused on a panel of 16 compounds: aspirin^[9]^, the angiotensin-converting enzyme (ACE) inhibitor captopril^[10]^, carbonyl cyanide m-chlorophenyl hydrazone (CCCP)^[11]^, the dual mammalian target of rapamycin (mTOR)/ Phosphoinositide 3-kinase inhibitor (PI3K) inhibitor GSK2126458, PI3K inhibitor LY-294002^[12]^, metformin^[13]^, antibiotics doxycycline^[14]^, minocycline^[15]^, and rifampicin^[16]^, nordihydroguaiaretic acid (NDGA)^[17]^, mTOR inhibitor rapamycin^[18]^, the statin simvastatin^[19]^, and natural compounds glycine^[20]^, caffeine^[21]^, resveratrol^[22]^ and urolithin A^[23]^.

Since most of the compounds that we tested were diluted in DMSO, we first tested the impact of DMSO concentration on *C. elegans* N2 lifespan with a live *E. coli* OP50 diet. Notably, increasing concentrations of DMSO up to 0.5% led to a slight, but significant, change in control median lifespan (15.2 vs. 15.8 days for H2O vs. 0.5% DMSO vehicle; **Figure S1**). DMSO concentrations from 1% to 2% more dramatically extended N2 lifespan (median >20 days) and was toxic at ≥5% DMSO (**Figure S1**). Subsequently, all lifespan experiments were carried out with up to 0.5% DMSO vehicle. We tested each compound for its ability to extend N2 survival when supplemented in the agar starting at the L4 stage under standard growth conditions with a live *E. coli* OP50 diet. Importantly, resveratrol (1mM; ΔMed = 85%), rifampicin (50uM; ΔMed = 59%), captopril (10mM; ΔMed = 48%), metformin (50mM; ΔMed = 40%), GSK2126458 (10uM; ΔMed = 36%), urolithin A (50uM; ΔMed = 35%), LY-294002 (50uM; ΔMed = 30%), CCCP (10uM; ΔMed = 29%), minocycline (100uM; ΔMed = 29%), doxycycline (10uM; ΔMed = 25%), and caffeine (5mM; ΔMed = 24%) significantly extended N2 lifespan at concentrations previously reported in the literature, further validating previous findings (**Figure 1A**).

**Figure 1.**
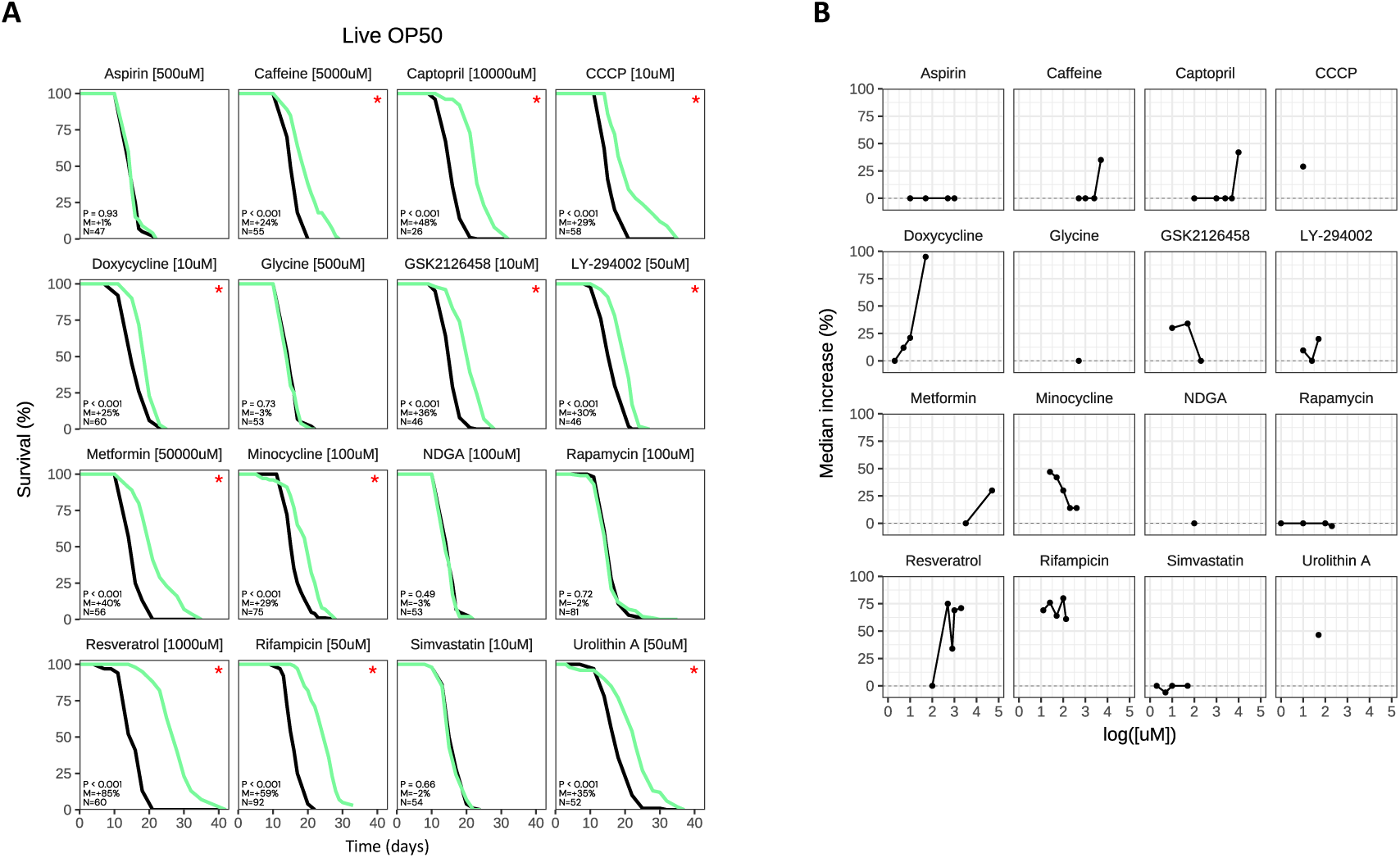
Effect on lifespan of 16 drugs in *C. elegans*, fed live OP50. (**A**) Survival curves for N2 treated with “Drug [Concentration]”, fed live OP50. P: p-value in the CoxPH model. M: median lifespan extension. N: sample size of the treated group. “*”: p < 0.05. Treated group in green, control group in black. Time in days from hatching. Representative result. (**B**) Dose response for each drug. Median lifespan increase (p < 0.05) in y-axis, log of concentration in x-axis in uM. Each dot represents a drug-concentration pair. Median extension = 0% if insignificant (p ≥ 0.05).

We further tested several of these compounds at multiple concentrations to determine if there was a dose-response effect. Indeed, caffeine and captopril did not extend lifespan at lower concentrations (**Figure 1B**) and minocycline had a reduced effect at higher concentrations (**Figure 1B**). On the other hand, doxycycline extended lifespan at relatively low concentrations, up to and including 50uM (**Figure 1B**). GSK2126458 best extended N2 lifespan when tested at 10uM, still retained a positive effect when tested at 50uM, and was no longer able to extend lifespan at 200uM (**Figure 1B**). Resveratrol extended lifespan at high concentrations (≥500uM) up to 2mM, and rifampicin extended lifespan at all tested concentrations (**Figure 1B**). Surprisingly, 5/16 compounds in our literature-curated panel did not extend lifespan in our system. Namely, neither aspirin (500uM), glycine (500uM), NDGA (100uM), rapamycin (100uM), nor simvastatin (10uM) significantly impacted N2 survival (**Figure 1A**). Moreover, testing several of these compounds at multiple concentrations did not yield any positive results (**Figure 1B**). Overall, these data further support resveratrol, rifampicin, urolithin A, captopril, caffeine, metformin, CCCP, minocycline, doxycycline, and LY-294002 as lifespan-extending compounds in *C. elegans*. Conversely, our data suggest that aspirin, glycine, NDGA, rapamycin, and simvastatin do not extend lifespan of *C. elegans* fed live OP50.

### Resveratrol, caffeine, metformin, doxycycline, GSK2126458, and LY-294002 extend C. elegans lifespan fed a UV-killed OP50 diet

Previous studies have found that the standard *C. elegans* diet, live *E. coli* strain OP50, could have a deleterious effect on lifespan due to the pathogenicity of the bacteria, potentially by causing pharyngeal swelling^[24]^. Additionally, it has been reported that some compounds may interact with the live bacteria, which could be responsible for their effect on lifespan^[13]^. To avoid these confounding effects, we decided to additionally perform lifespan experiments on N2 fed UV-killed OP50. First, we determined the impact of UV-killed OP50 on N2 lifespan. As previously observed^[25]^, *C. elegans* on a UV-killed OP50 diet lived significantly longer than those on a live OP50 diet (ΔMed = 42-48%), supporting the argument that live OP50 is pathogenic to *C. elegans* (**Figure 2A; Figure S1**).

**Figure 2.**
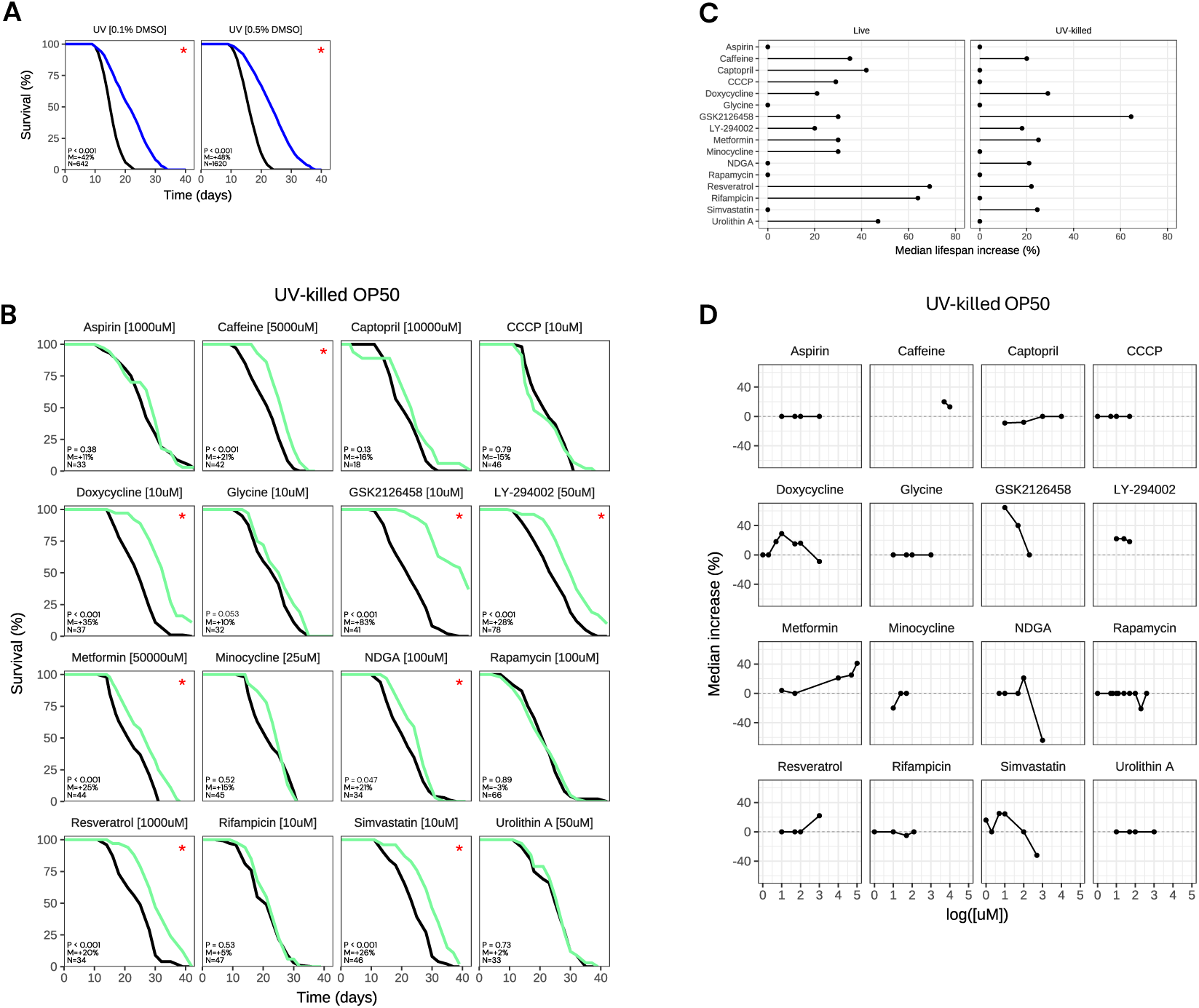
Resveratrol, caffeine, metformin, doxycycline, GSK2126458, and LY-294002 still extend *C. elegans* lifespan, fed a UV-killed OP50 diet. (**A**) Effect of UV-killed OP50 on wild-type N2 lifespan, with exposure to 0.1% or 0.5% DMSO vehicle. (**B**) Survival curves for N2 treated with “Drug [Concentration]”, fed UV-killed OP50. P: p-value in the CoxPH model. M: median lifespan extension. N: sample size of the treated group. “*”: p < 0.05. Treated group in green, control group in black. Time in days from hatching. Representative result. (**C**) Summary of the effect of 16 drugs in live and UV-killed OP50 on N2 median lifespan, at the best concentration tested. Median extension = 0% if insignificant (p ≥ 0.05). (**D**) Dose response for each drug. Median lifespan increase (p < 0.05) in y-axis, log of concentration in x-axis in uM. Each dot represents a drug-concentration pair. Median extension = 0% if insignificant (p ≥ 0.05).

Here again, we first tested the impact of DMSO concentration on N2 lifespan fed with UV-killed OP50. Notably, we found that the percentage of DMSO in the agar impacted *C. elegans* lifespan. Namely, DMSO concentrations up to 2% increasingly extended N2 lifespan and was toxic at ≥5% DMSO (**Figure S1**). For this reason, we sought to keep the concentration to ≤0.2% and only up to 0.5% in necessary cases. We next tested our literature-curated panel of 16 compounds at multiple concentrations in *C. elegans* fed UV-killed OP50 (**Figure 2B-D**). Of the 11 compounds that extended survival with live OP50, 6 compounds still reproducibly extended lifespan when fed UV-killed OP50, including resveratrol (1mM; ΔMed = 20%), metformin (50mM; ΔMed = 25%), GSK2126458 (10uM; ΔMed = 83%), LY-294002 (50uM; ΔMed = 28%), doxycycline (10uM; ΔMed = 35%), and caffeine (5mM; ΔMed = 21%). Interestingly, resveratrol and metformin extended lifespan to a lesser effect when fed UV-killed OP50 than live OP50. Conversely, GSK2126458, and to a lesser extent doxycycline, had greater impact on median lifespan when fed UV-killed OP50. Overall, these six compounds were able to extend *C. elegans* lifespan with both live and UV-killed OP50. Separately, of the 5 compounds that did not extend N2 lifespan when fed live OP50, 3 of them continued to fail to extend lifespan when tested at ≥4 concentrations (aspirin, glycine, and rapamycin) when fed UV-killed OP50 (**Figure 2D**). Conversely, the other two compounds NDGA and simvastatin extended lifespan with UV-killed OP50 at one concentration at least (**Figure 2D**), suggesting that their effect may only be seen in that context. Altogether, these results demonstrate that a live or UV-killed OP50 diet can impact the potential lifespan-extending effect of a compound in *C. elegans*.

### Direct effect of the compounds on OP50 viability

Of the 5 compounds that extended lifespan with live OP50 but not with UV-killed OP50, 3 of them (rifampicin, minocycline, and CCCP) are known to decrease live *E. coli* viability. Given our finding that UV-killed OP50 increases N2 survival, likely due, at least in part, to pathogenicity of live *E. coli*, this raises the possibility that these three compounds increased lifespan when fed live OP50 by directly killing the bacteria rather than having an effect on the worm itself. To confirm this hypothesis, we sought to directly assess the effect of positive hits from the live OP50 screen on the bacteria alone. To accomplish this, we incubated each compound with seven 1:10 serial dilutions of live OP50 in a 96-well plate containing LB media overnight at 37°C and observed OP50 growth the next morning.

Out of the 11 compounds that extended lifespan with live OP50 diet, 7 directly negatively affected OP50. We first confirmed that rifampicin, minocycline, and CCCP did indeed decrease live OP50 viability, however to different degrees – minocycline completely killed OP50 regardless of concentration, rifampicin only after the first 1:10 dilution, and CCCP only decreased OP50 thickness at each dilution, instead of completely killing it (**Table 1, Figure S2**). Notably, doxycycline entirely killed OP50 regardless of OP50 concentration, similarly to minocycline, which is unsurprising given that they are known antibiotics. More surprisingly, we found that captopril, metformin, and to a lesser extent caffeine, also decreased OP50 viability (**Table 1, Figure S2**).

**Table 1.** Direct effect of the drug on OP50 viability. Direct effect of the drug-concentration tested on the OP50 alone. Summary of the effect of 16 drugs in live and UV-killed OP50 on N2 median lifespan. The impact of the drug on OP50 was determined by the percentage of wells with viable OP50 at the multiple OP50 dilutions tested (7 1:10 serial dilutions starting from 0.3mg/mL OP50) and the quality of the OP50 pellets. Quality of OP50 pellets was determined by visual assessment (Figure S2).

| Drug | Conc. (uM) | ΔMedian Lifespan with Live OP50 | ΔMedian Lifespan with UV-killed OP50 | Drug Negatively Affects OP50? | % Wells with OP50 | Overall quality of OP50 pellet |
| --- | --- | --- | --- | --- | --- | --- |
| Resveratrol | 1000 | 82% | 20% | No | 100% | Normal |
| Rifampicin | 10 | 69% | 0% | Yes | 14% | Small |
| Urolithin A | 50 | 56% | 0% | No | 100% | Normal |
| Captopril | 10000 | 53% | 0% | Yes | 14% | Sparse |
| CCCP | 10 | 45% | 0% | Yes | 100% | Sparse |
| Caffeine | 5000 | 44% | 21% | Yes | 100% | Sparse |
| Metformin | 50000 | 37% | 25% | Yes | 29% | Sparse |
| Minocycline | 25 | 37% | 0% | Yes | 0% | N/A |
| LY-294002 | 50 | 24% | 28% | No | 100% | Normal |
| Doxycycline | 10 | 23% | 35% | Yes | 0% | N/A |
| Simvastatin | 10 | 0% | 26% | No | 100% | Normal |

Importantly, resveratrol, urolithin A and LY-294002 did not affect the OP50, suggesting that their lifespan-extending effect cannot be attributed to decreasing bacteria viability. Additionally, resveratrol and LY-294002 also extended lifespan of worms fed with UV-killed OP50. On the other hand, rifampicin, captopril, CCCP, minocycline extended lifespan of *C. elegans* only when fed live OP50, and directly reduced OP50 viability, suggesting that their lifespan-extending effect may be due to a negative impact of the compound on OP50. Caffeine, metformin, and doxycycline affected E. coli viability, but also extended lifespan when fed UV-killed OP50, therefore their positive effect on lifespan cannot be explained by their effect on the bacteria alone. Taken together, these results demonstrate that the effect of compounds on lifespan when fed live *E. coli* cannot be entirely explained by their effect on the bacteria.

### Combination studies identify beneficial drug pairs

After confirming which of the 16 compounds robustly extended *C. elegans* lifespan, we next sought to determine if additive effects could be observed by combining some of the positive hits from our initial screen. Indeed, combinatorial interventions hold the potential to increase effect sizes, by targeting multiple aging pathways at the same time. More precisely, we tested pairwise combinations of four of our best hits, namely doxycycline (10uM), LY-294002 (50uM), GSK2126458 (10uM), and resveratrol (1mM); both with live and UV-killed OP50 diets. We found that many combinations did not yield any significant increase in lifespan extension, compared to the single compounds alone (13 out of 16 combinations, **Figure 3A-B**). Surprisingly, some combinations even cancelled the positive effect of the single compounds alone. Combinations of resveratrol with GSK2126458 or LY-294002 were largely inferior to resveratrol alone when fed live OP50. With a UV-killed diet, the combination of resveratrol with GSK2126458 was inferior to both resveratrol and GSK2126458 alone, even reaching slight toxicity. On the other hand, combination of GSK2126458 with doxycycline or LY-294002 dramatically increased the effect of each single drug alone with a live OP50 diet (p < 0.001 or p = 0.01, respectively, for the combination compared to best single compound). With a UV-killed diet, combining doxycycline and resveratrol also had an increased effect compared to single drugs alone (p = 0.0413, for the combination compared to best single compound, **Figure 3A-B**).

**Figure 3.**
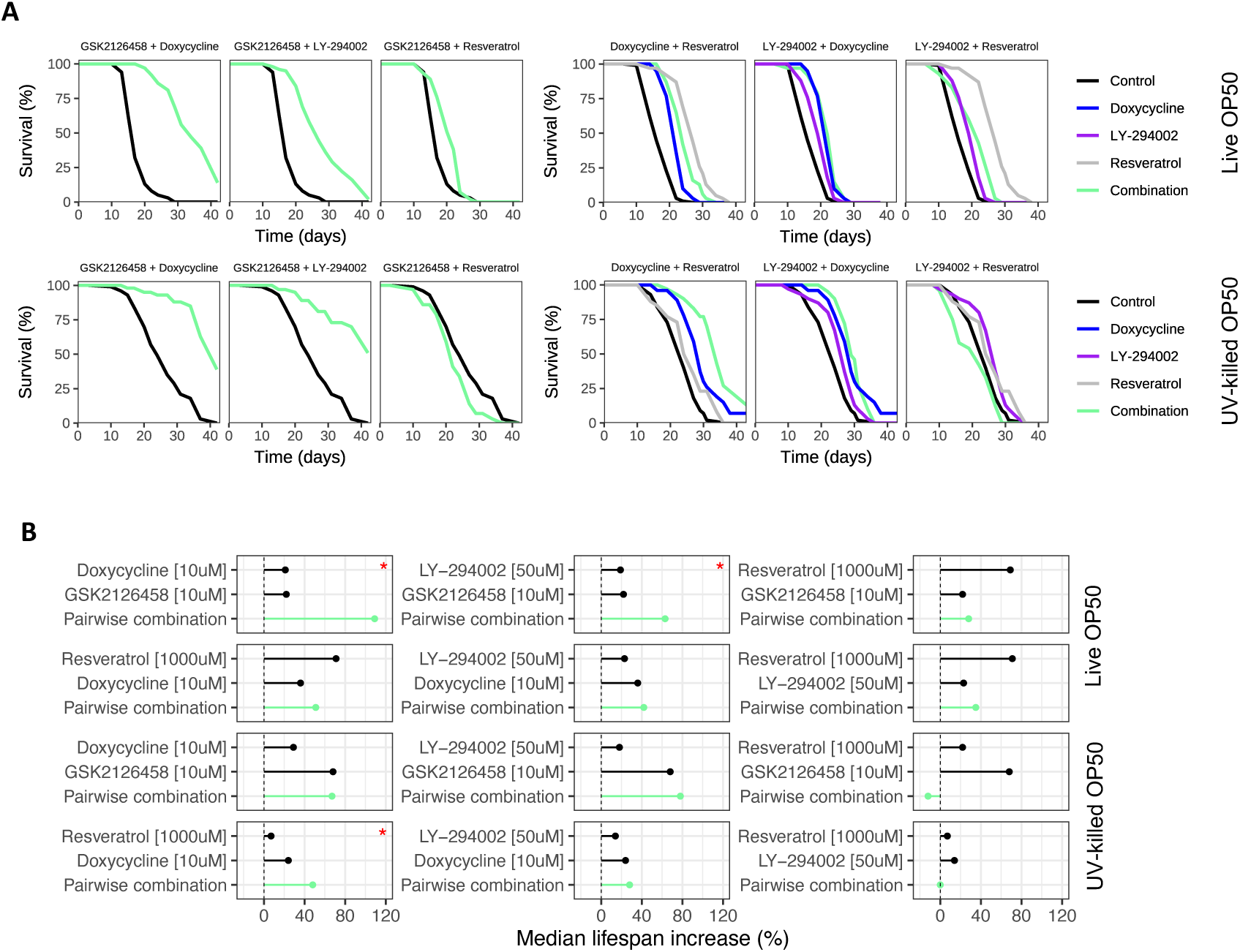
Combination studies identify beneficial drug pairs. (**A**) Survival curves for pairwise combinations of drugs tested on N2 lifespan when fed live OP50. Control in black, single drugs in blue, grey, purple, pairwise combination in light green. “*”: p < 0.05 for comparison of combination survival to all the single-drug results. (**B**) Summary effect of the single drugs and combinations on N2 lifespan from the survival curves. Median extension of the single compounds for comparison to the GSK2126458 combinations is a representative result. Median extension = 0% if insignificant (p ≥ 0.05). “*” indicates combination result is significant compared to each single drug alone.

Lastly, we also tested pairwise combinations of two other drug triplets: minocycline (100uM), caffeine (5mM), and rifampicin (50uM), with a live OP50 diet; and LY-294002 (50uM), caffeine (5mM), and metformin (50mM), with a UV-killed OP50 diet. 5 out of 6 combinations did not yield any significant beneficial effect, compared to single molecules alone. However, the combination of rifampicin and caffeine had a nearly additive impact on median lifespan (p < 0.001, for the combination compared to best single compound, **Figure S3A-B**). Taken altogether, these results demonstrate a relatively low hit rate for combination experiments but suggest a possible benefit in the combination of GSK2126458 with LY-294002 or doxycycline and argue against combinations of resveratrol with LY-294002 or GSK2126458.

### Impact of resveratrol and LY-294002 on lifespan of other Caenorhabditis nematodes

As the effect of molecules in *C. elegans* has been reported to be strain-dependent^[5]^, we sought to test the effect of two of our best compounds in multiple nematode strains. For this reason, we tested the effect of LY-294002 and resveratrol in *C. elegans* (N2, JU775), as well as *C. tropicalis* (JU1373), and *C. briggsae* (AF16) with both live and UV-killed OP50 diets. Interestingly, LY-294002 (50uM) significantly extended JU775 lifespan when fed live OP50 (ΔMed = 29%) and UV-killed OP50 (ΔMed = 21%). Resveratrol (1mM) extended JU775 lifespan with live OP50 (ΔMed = 84%) but did not impact lifespan when fed UV-killed OP50 (**Figures 4A-B**). We found that LY-294002 (50uM) and resveratrol (1mM) did not impact JU1373 when fed live OP50 (**Figure 4A-B**), but both compounds significantly increased JU1373 median lifespan when fed UV-killed OP50 (ΔMed = 45% or 65%, respectively; **Figure 4A-B**). Interestingly, we found that AF16 did not tolerate the UV-killed OP50 diet, and largely fled wells containing this diet (data not shown). Additionally, both LY-294002 (50uM) and resveratrol (1mM) significantly extended AF16 lifespan (ΔMed = 13% or 51%, respectively; **Figure 4A-B**). Overall, the lifespan-extending effect of resveratrol, and more strongly LY-294002, was conserved across most of the conditions tested in these different strains and diets.

**Figure 4.**
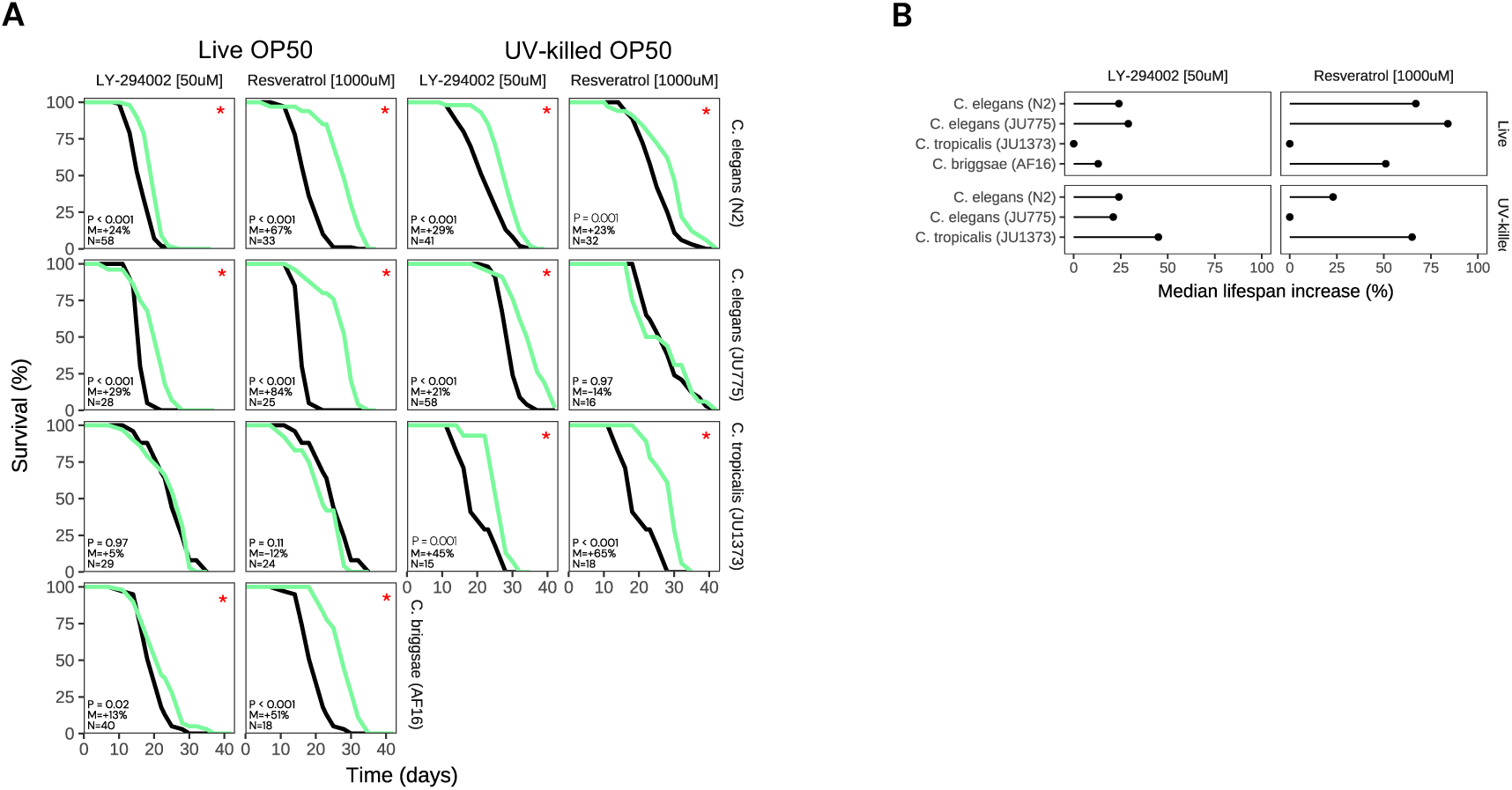
Impact of resveratrol and LY-294002 on lifespan of other Caenorhabditis nematodes. (**A**) Survival curves for *C. elegans* (N2 or JU775), *C. tropicalis* (JU1373), and *C. briggsae* (AF16) treated with LY-294002 (50uM) or resveratrol (1000uM), fed either live or UV-killed OP50. P: p-value in the CoxPH model. M: median lifespan extension. N: sample size of the treated group. “*”: p < 0.05. Treated group in green, control group in black. Time in days from hatching. (**B**) Summary effect of LY-294002 (50uM) or resveratrol (1000uM) on Caenorhabditis change in median survival relative to vehicle control. Median extension = 0% if insignificant (p ≥ 0.05).

### Doxycycline, caffeine, and GSK2126458 extend lifespan when administered at day 1 of adulthood

When evaluating the effect of a compounds on the lifespan of organisms, the starting time of treatment is critical. While some compounds in the literature have been shown to be able to extend lifespan only when given during development, others are able to retain their effect even when given at a later stage in life^[26,27]^. In this study, we exposed worms to the compounds for their entire lifespan starting at the late L4 stage, the final larval stage. The reason for this protocol is that this also coincides with the strict timing of FUdR, a chemical that suppresses progeny formation, which is used in most lifespan experiments. Thus, we wondered if by delaying compound treatment by just one day, allowing worms to fully reach adulthood, we would see a difference in the effect of the compounds on lifespan. Thus, we transferred day 1 adult N2 *C. elegans* to experimental plates containing a subset of our best positive compounds, fed a diet of live or UV-killed OP50.

Notably, we found that doxycycline extended *C. elegans* lifespan when administered at L4 and day 1 adult stages, regardless of diet and concentration tested (**Figure 5A-D, Supplemental Table 1**). Moreover, caffeine (5mM), metformin (50mM), resveratrol (800uM), rifampicin (50uM), and urolithin A (50uM), still significantly extended N2 survival when administered at day 1 of adulthood and fed live OP50, however median extension was less than what we typically observed when administered at L4 (**Figure 5A-B**, **Supplemental Table 1**). Interestingly, LY-294002 (50uM) and GSK2126458 (10uM), did not extend N2 lifespan when administered at day 1 and fed live OP50 (**Figure 5A-B**). When fed UV-killed OP50, caffeine (5mM) and simvastatin (10uM) still extended lifespan when administered at day 1 of adulthood, with only a slight decrease in median extension compared to L4 (**Figure 5C-D**). Metformin was no longer able to extend lifespan when fed UV-killed OP50 (**Figure 5C-D**). Importantly, GSK2126458 (10uM) greatly extended lifespan when administered at day 1 of adulthood when fed UV-killed OP50 (ΔMed = 68%), comparable to its effect at the L4 stage, while LY-294002 (50uM) was also not able to extend lifespan at this stage (**Figure 5C-D**). Overall, most of the compounds evaluated extend lifespan when administered at day 1 of adulthood, although sometimes at lower potency when tested at the same concentration as the L4 experiments.

**Figure 5.**
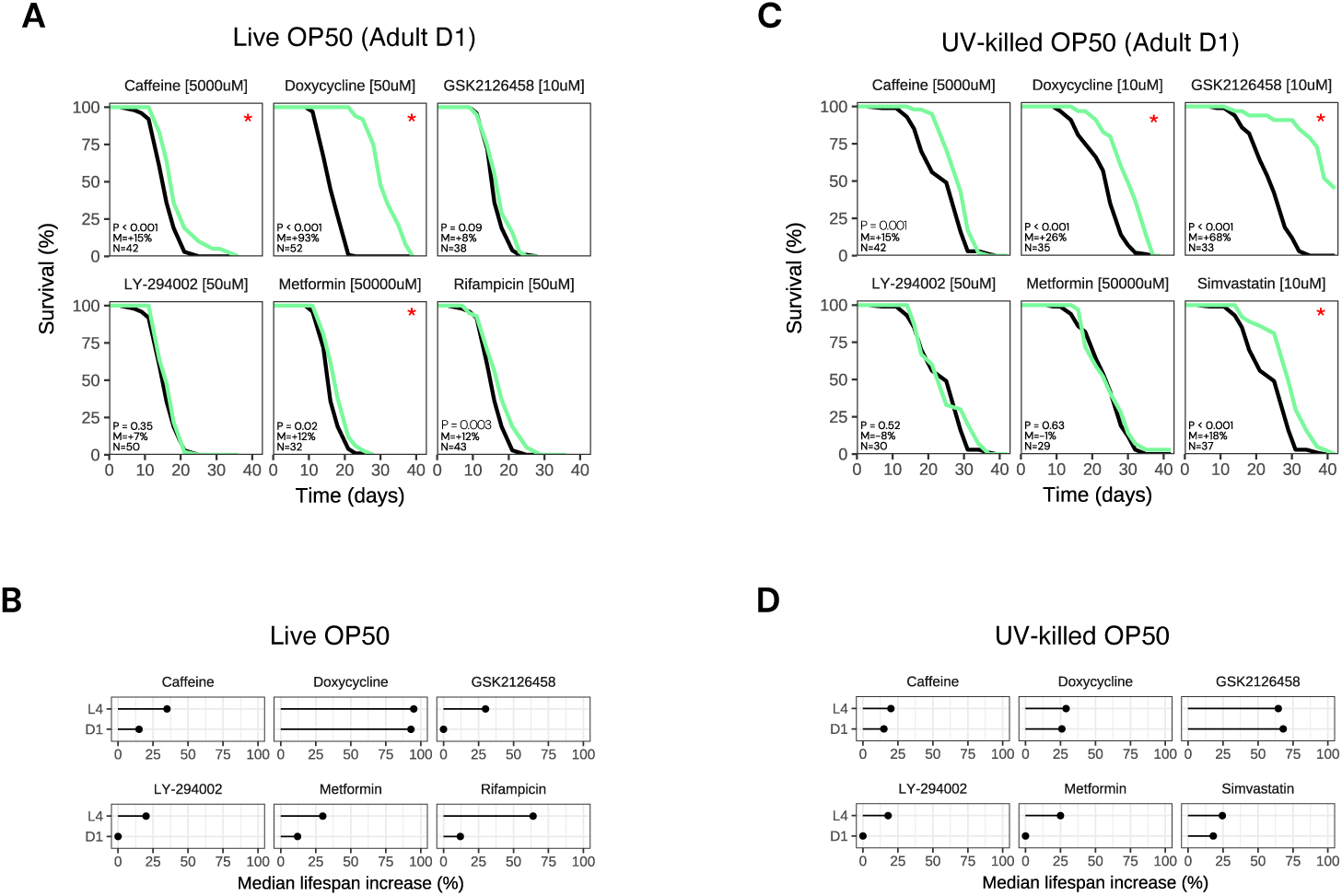
Doxycycline and caffeine extend lifespan when administered at day 1 of adulthood with both a live and UV-killed OP50 diet. (**A**) Survival curves for N2 treated with “Drug [Concentration]” starting at day 1 of adulthood, fed live OP50. P: p-value in the CoxPH model. M: median lifespan extension. N: sample size of the treated group. “*”: p < 0.05. Treated group in green, control group in black. Time in days from hatching. (**B**) Summary effect of each drug on median lifespan extension when administered at late L4 versus day 1 of adulthood, fed live OP50. Median extension = 0% if insignificant (p ≥ 0.05). (**C**) Survival curves for N2 treated with “Drug [Concentration]” starting at day 1 of adulthood, fed UV-killed OP50. P: p-value in the CoxPH model. M: median lifespan extension. N: sample size of the treated group. “*”: p < 0.05. Treated group in green, control group in black. Time in days from hatching. (**D**) Summary effect of each drug on median lifespan extension when administered at late L4 versus day 1 of adulthood, fed UV-killed OP50. Median extension = 0% if insignificant (p ≥ 0.05).

### Doxycycline, metformin, rifampicin, and LY-294002 extend D. melanogaster lifespan

We next asked whether the lifespan extending effect we observed in *C. elegans* could be conserved across species. For this reason, we tested a subset of the *C. elegans* positive molecules in a larger model organism, the fly *D. melanogaster*. Towards this goal, we incorporated doxycycline, metformin, rifampicin, LY-294002, or minocycline into the fly diet, and measured their impact on *D. melanogaster* lifespan. Interestingly, we found that doxycycline (500uM) significantly extended both male and female fly lifespan (ΔMed = 34% and 29%, respectively), and extended lifespan at lower concentrations (**Figure 6**). Additionally, metformin (15mM) increased male and female lifespan (Δmed = 16% and 27%, respectively; **Figure 6**). Similarly, rifampicin (50uM) increased male and female lifespan (ΔMed = 18% and 18%, respectively; **Figure 6**). LY-294002 (50uM) led to lifespan extension in male (ΔMed = 13%) but not female flies (**Figure 6**). Conversely, minocycline (200uM) did not extend neither male nor female lifespan. Out of the 5 compounds that we tested 4 had a conserved effect on *D. melanogaster* lifespan extension, and one did not.

**Figure 6.**
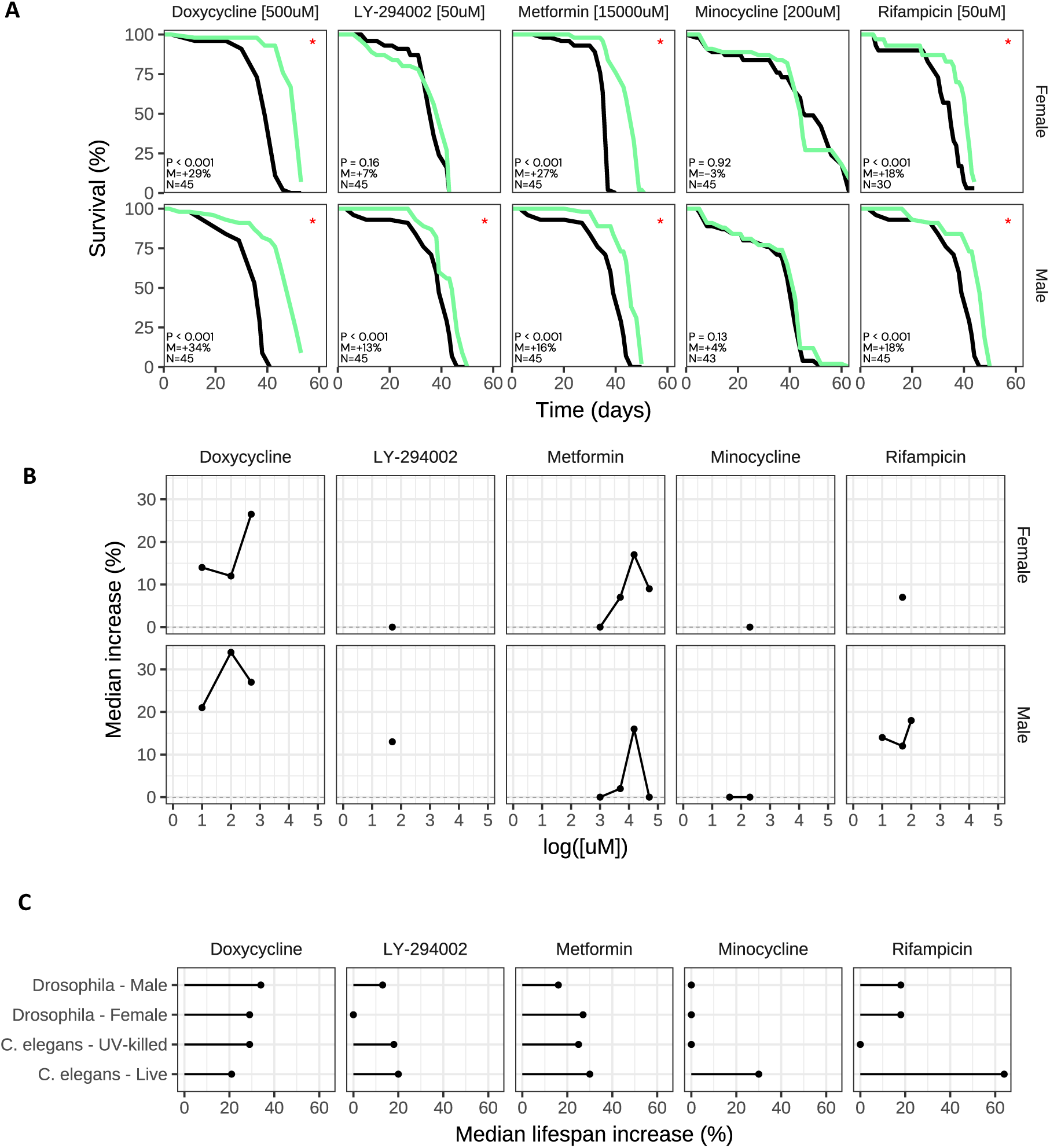
Doxycycline, metformin, rifampicin, and LY-294002 extend *D. melanogaster* lifespan. (**A**) Survival curves for Canton S male or female *Drosophila melanogaster* treated with “Drug [Concentration]”. P: p-value in the CoxPH model. M: median lifespan extension. N: sample size of the treated group. “*”: p < 0.05. Treated group in green, control group in black. Representative result. **(B)** Dose response for each drug. Median lifespan increase (p < 0.05) in y-axis, log of concentration in x-axis in uM. Median extension = 0% if insignificant (p ≥ 0.05). (**C**) Summary effect of each drug on median lifespan extension for male or female *Drosophila* compared to N2 *C. elegans* fed either live or UV-killed OP50. Median extension = 0% if insignificant (p ≥ 0.05).

### Large-scale screening of putative lifespan extending compounds in C. elegans identifies miconazole, fenbendazole, tadalafil, lotilaner, and safinamide as novel candidates

We next expanded our study to screen more than 200 compounds for extension of N2 lifespan when administered at the late L4 stage and fed either live (195 compounds) or UV-killed (228 compounds) OP50. Each compound was tested at a minimum of two different concentrations. Full survival results are included in **Supplemental Table 1**. This small library contained compounds from different categories, such as antioxidants, anti-inflammatories, antibiotics and anti-fungals, PI3K inhibitors, anti-diabetics, hormones, psychotropics, and antihypertensives. Overall, of all the compounds and concentrations tested, 34/470 (7.2%) of experiments with live OP50 and 39/592 (6.7%) of experiments with UV-killed OP50 yielded a significantly (p<0.05) positive median lifespan extension (**Figure 7A, Supplemental Table 1**). We also observed a low variability within our control conditions, when fed either live or UV-killed OP50, suggesting high consistency (**Figure S4**).

**Figure 7.**
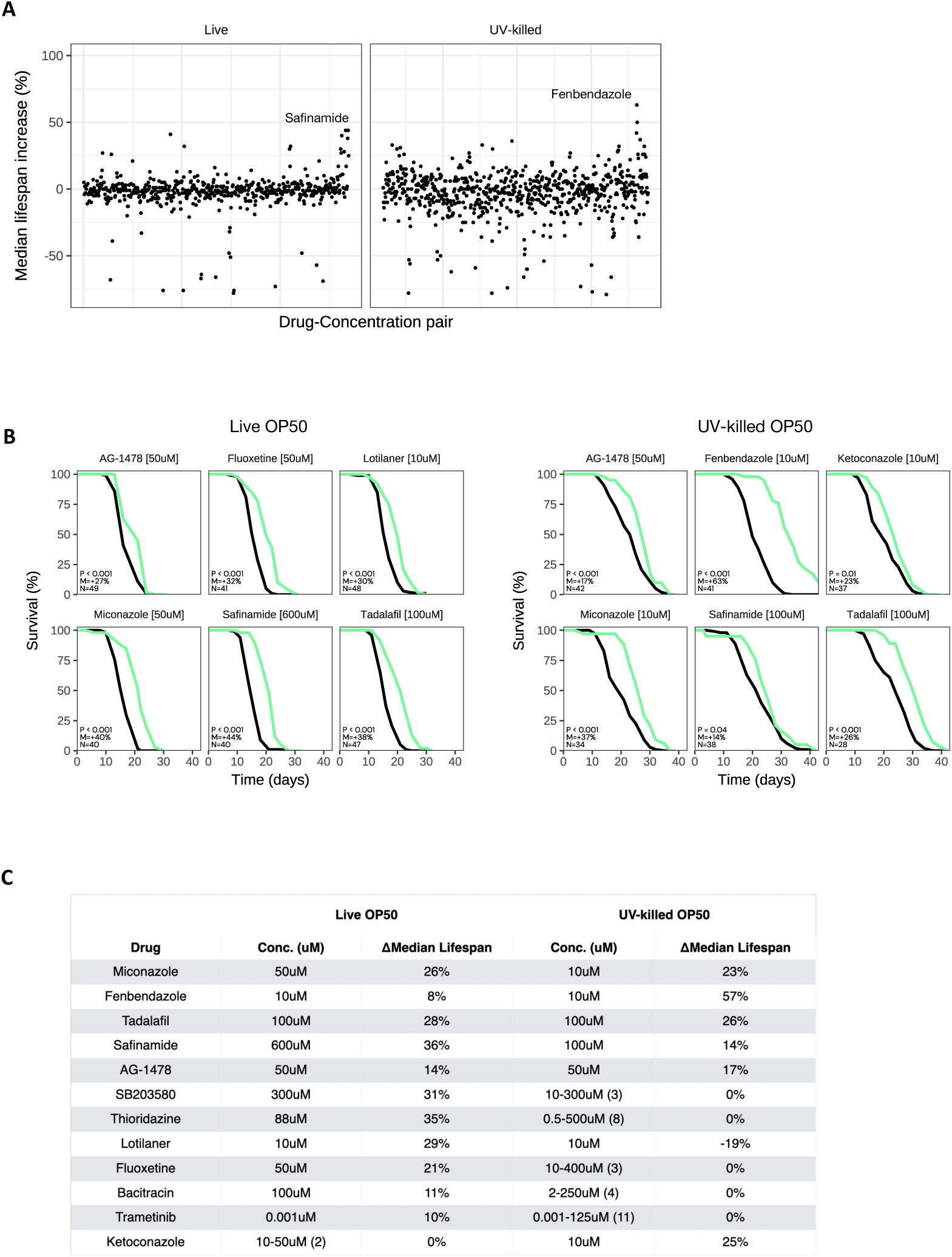
Large-scale screening of putative lifespan extending compounds in *C. elegans*. (**A**) Dot-plot of the impact of ≥200 compounds (≥2 concentrations tested) on N2 median lifespan when fed live (left panel) or UV-killed (right panel) OP50. Each dot represents and individual experiment. Median lifespan increase (%) in y-axis and drugs in alphabetical order in x-axis. (**B**) Survival curves for top lifespan-extending drugs when N2 are fed live (left panel) or UV-killed (right panel) OP50. (**C**) Summary of top survival results. Median extension = 0% if insignificant (p ≥ 0.05). The range of concentrations tested are listed with total number of concentrations tested in “()” for drugs that did not significantly change median lifespan.

We repeated the top hits from the screen and ultimately identified 11 compounds that reproducibly extended lifespan when fed live OP50 and 6 with UV-killed OP50 (**Figure 7B-C**). 5 of these compounds extended lifespan when fed both live and UV-killed OP50: antifungal agent miconazole (ΔMed = 40% and 37%, respectively), antiparasitic fenbendazole (ΔMed = 8% and 63%, respectively), PDE5 inhibitor tadalafil (ΔMed = 38% and 26%, respectively), MAO-B inhibitor safinamide (ΔMed = 44% and 14%, respectively), and tyrosine kinase inhibitor AG-1478 (ΔMed = 27% and 17%, respectively; **Figure 7B-C**). Of these 5, only AG-1478 had been previously tested and shown to extend lifespan in *C. elegans*^[28]^.

Of the 11 compounds found to reproducibly extend lifespan in N2 when fed live OP50, 6 did not extend N2 lifespan when fed UV-killed OP50 at ≥3 concentrations tested. Namely, p38 MAPK inhibitor adezmapimod (SB203580) (ΔMed = 32%), antibiotic bacitracin (ΔMed = 11%), antidepressant fluoxetine (ΔMed = 32%), anti-parasitic lotilaner (ΔMed = 30%), and antipsychotic thioridazine (ΔMed = 25%) extended lifespan only when fed live OP50 (**Figure 7B**). Of these compounds, thioridazine, fluoxetine, and bacitracin have previously been tested and shown to extend lifespan in *C. elegans*^[14,29,30]^, and adezmapimod has been shown to extend lifespan in *D. melanogaster*^[31]^. Additionally, we identified one compound that extended *C. elegans* lifespan when fed UV-killed OP50 but not live OP50 when tested at ≥3 concentrations, the antifungal ketoconazole (ΔMed = 23%, **Figure 7B**).

We thus identify miconazole, fenbendazole, tadalafil, safinamide, adezmapimod (SB203580), lotilaner, and ketoconazole as novel lifespan extending compounds in *C. elegans*. We also validate the lifespan extending effects of thioridazine, fluoxetine, and bacitracin, dependent on OP50 diet.

## Discussion

In this study, we sought to validate a panel of 16 compounds reported to extend lifespan in *C. elegans*. We found that 11 compounds, including resveratrol, rifampicin, captopril, metformin, GSK2126458, and urolithin A, significantly extended lifespan when fed a live *E. coli* OP50 diet. We identified dose-response effects, particularly for caffeine and captopril, which required higher doses, while doxycycline and rifampicin were effective across a range of concentrations. We confirmed that a UV-killed OP50 diet extends *C. elegans* lifespan, and revealed that only six compounds (resveratrol, metformin, GSK2126458, LY-294002, doxycycline, and caffeine) retained their lifespan-extending effects with this diet. The reduced efficacy of rifampicin, captopril, CCCP, and minocycline with this diet suggests that their effect with live OP50 may be due their antibiotic activity rather than a direct effect on *C. elegans*. This was further confirmed by assessing their impact on OP50 viability, where these compounds significantly reduced bacterial growth.

Combination studies identified beneficial drug pairs, notably rifampicin and caffeine, GSK2126458 and doxycycline, and GSK2126458 and LY-294002, which together produced more than additive effects on lifespan extension. Conversely, combining resveratrol with LY-294002 or GSK2126458 abolished their individual positive effects. In our study, combining two compounds with positive effect on lifespan rarely led to an additive or synergistic effect and sometimes canceled the positive effect observed in the single compounds. These results highlight the importance of combinatorial screenings for the identification of combinations of compounds with synergistic effects of lifespan, and strongly suggest against the consumption of multiple supplements and drugs, which combined effects on health and lifespan have yet to be evaluated.

Moreover, we showed that resveratrol and LY-294002 maintained their lifespan-extending effects across various nematode strains and diets, with some strain-specific differences. Furthermore, delayed compound administration, at day 1 of adulthood, generally retained efficacy, albeit sometimes with reduced potency. Using *D. melanogaster*, we also show that the effect of some compounds on lifespan can be conserved across species, while others may be species dependent, increasing the potential translation of these discoveries to larger animals including companion animals and humans. Lastly, in a small proof-of-concept screening of ∼200 compounds, we identified new compounds extending lifespan in *C. elegans*.

In this study we highlight the complex relationship between lifespan-extending compounds and the *C. elegans* diet. We find examples of compounds that extend lifespan with a live OP50 diet and toxic to the bacteria; that extend lifespan with a live OP50 diet and not toxic to the bacteria; and that extend lifespan with a live OP50 diet, toxic to the bacteria, but also extending lifespan with UV-killed OP50. Based on these results, it is difficult to conclude on the role of the interactions between compounds and the bacteria on *C. elegans* lifespan extension. Further studies will be needed to increase our understanding of this important aspect. Until then, we recommend testing compounds with both live and UV-killed, and separately assessing the effect of the compounds on bacteria alone.

Previous studies have found that *C. elegans* lifespan, and the effect of putative lifespan-extending molecules, can vary due to even subtle changes in the environment, compound manufacturers, and experimental conditions^[32,33]^. For example, rapamycin, a popular lifespan-extending drug candidate, has been shown to extend lifespan across different species including *C. elegans*^[18,34]^. However, other groups, including the CITP, which specifically tests compounds in *C. elegans* in three different sites to thoroughly interrogate a candidate, have similarly not reproduced lifespan extension by rapamycin in *C. elegans*^[5]^. Other groups have reported recently that the reason may be that rapamycin precipitates out of the agar, something we have also observed, depending on concentrations.

Overall, our study demonstrates that while several compounds can extend lifespan in *C. elegans*, their efficacy is influenced by bacterial diet, dosage, strain specificity, and starting age of the treatment. Our results underscore the importance of validating lifespan-extending interventions under standardized conditions and exploring potential combinations for enhanced efficacy.

## Methods

### *C. elegans* maintenance

Worms [N2, JU775, JU1373, AF16] were obtained from the Caenorhabditis Genetics Center (CGC). Worms were maintained at 20°C and 80% humidity in standard solid NGM growth conditions [0.3% NaCl, 0.25% BactoPeptone, 2% agar, 1 mM MgSO₄, 1 mM CaCl₂, 5 µg/mL cholesterol, and 25 mM potassium phosphate buffer (pH 6.0)]. Unless otherwise stated, worms were fed live OP50 obtained fresh from overnight inoculation of a single OP50 colony into LB at 37°C, concentrated to 30mg/mL in ddH2O, and seeded on solid NGM agar plates. OP50 was obtained from the CGC.

### Compounds

A complete list of the compounds tested, and catalog numbers is provided in **Supplemental Table 1**. All compounds were dissolved in H2O or DMSO dependent on solubility.

### Lifespan assay

Detailed and summarized data for all lifespan experiments carried out in this study can be found in **Supplemental Table 1**.

Worms were synchronized by 10-min hypochlorite treatment (1% sodium hypochlorite, 800mM sodium hydroxide, and water) and washed 4 times with M9 buffer to isolate embryos. Embryos were hatched overnight in M9 buffer on a rocker at 20°C (day = 0 on lifespan graphs). L1s were then transferred to NGM plates seeded with live OP50. Unless otherwise noted, worms were transferred to experimental plates at the late L4 stage (15 worms/well of a 24-well plate; 3 wells per condition). For experiments that exposed worms to drug at day 1 of adulthood, late L4 worms were transferred to maintenance plates containing 150uM 5-Fluoro-2’deoxyuridine (FUdR; calculated with respect to the volume of the agar to prevent progeny production). 24 hours later, day 1 adult worms were transferred to experimental plates.

24-well experimental plates were prepared by mixing drug or vehicle into NGM agar (1.5mL/well) prior to solidification. Drug concentration was calculated with respect to the volume of the agar and treated and control wells contained an equal concentration of DMSO. Experimental plates were seeded with 30uL of live or UV-killed OP50 (60mg/mL) mixed with 150uM FUdR (calculated with respect to the volume of the agar). UV-killed OP50 was prepared by exposing live OP50 diluted to 30mg/mL to 254nm high-intensity UV light for 4 hours on a UV transilluminator (TFS-30V) and subsequently concentrated to 60mg/mL before seeding on experimental NGM plates. Experimental plates were wrapped in parafilm and kept at 20°C at 80% humidity for the duration of the experiment. Worm lifespan was assessed by movement over 48-hour periods using standard microscopy.

### Survival analysis

Statistics for the survival analysis were obtained by fitting a proportional hazards regression model (Cox-PH) on treated compared to control worms. Median extension, p-value of the log-rank test and sample size were reported.

### Significance of combination assays

For combinatorial assays where we tested simultaneously single compounds and combinations at the same time, the combination was deemed significant if it was significant compared to the best single compound, in the sense of the log-rank test. For combinatorial assays where we did not have the single compounds tested as part of the same experiment as the combination, we assessed significance using Fisher’s Z-test.

### OP50 viability assay

A compound or vehicle were added at the desired concentration (max 0.5% DMSO) to a final volume of 100uL of liquid LB in a round-bottom 96-well plate, 7 wells each per condition. 1uL of 30mg/mL OP50 was added to the first experimental well per condition, and 6 1:10 serial dilutions of OP50 were subsequently carried out. Plates were incubated in a shaker at 37°C overnight (18hrs). The next day, OP50 was allowed to pellet, and plates were imaged to assess OP50 growth.

### Drosophila lifespan assay

Canton S *D. melanogaster* flies were obtained from the Bloomington Drosophila Stock Center (BDSC). 45 male and female flies were sorted upon hatching (virgin flies) and maintained at 25°C on a 12-h light/dark cycle, at constant 50% humidity. Expired animals were counted by visually assessing lack of movement inside the vial. The compound was mixed into a Cornmeal diet, supplemented with 0.05% ampicillin to prevent bacterial growth.

## Acknowledgements

We thank all the team members of EPITERNA.

## Author contribution

G.P. performed all the worm experiments and co-wrote the manuscript. J. M. performed data and statistical analysis. C. P. performed the worm and fly experiments. L. S. helped with the worm experiments. S. G. helped prepare the worm and fly experiments. A. O. co-supervised the work, designed the study, and co-wrote the manuscript. K. P. co-supervised the work, designed the study, performed data analysis and visualization, and co-wrote the manuscript.

## Data sharing

All data produced in the present work are contained in the manuscript. No additional data available.

---

**Figure S1.**
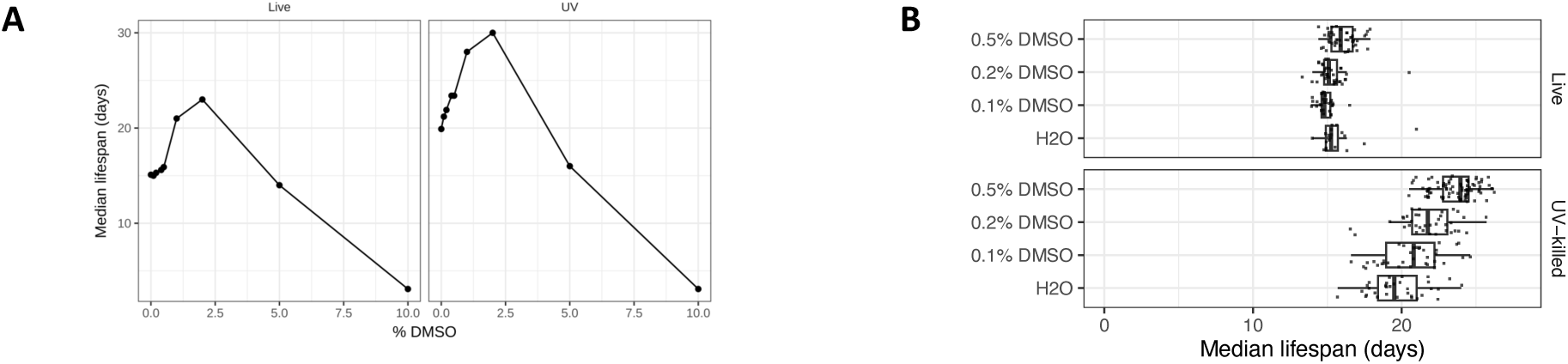
Impact of DMSO concentration on *C. elegans* survival. Percent of DMSO in NGM agar versus median lifespan of N2 *C. elegans* fed either live (left) or UV-killed (right) OP50.

**Figure S2.**
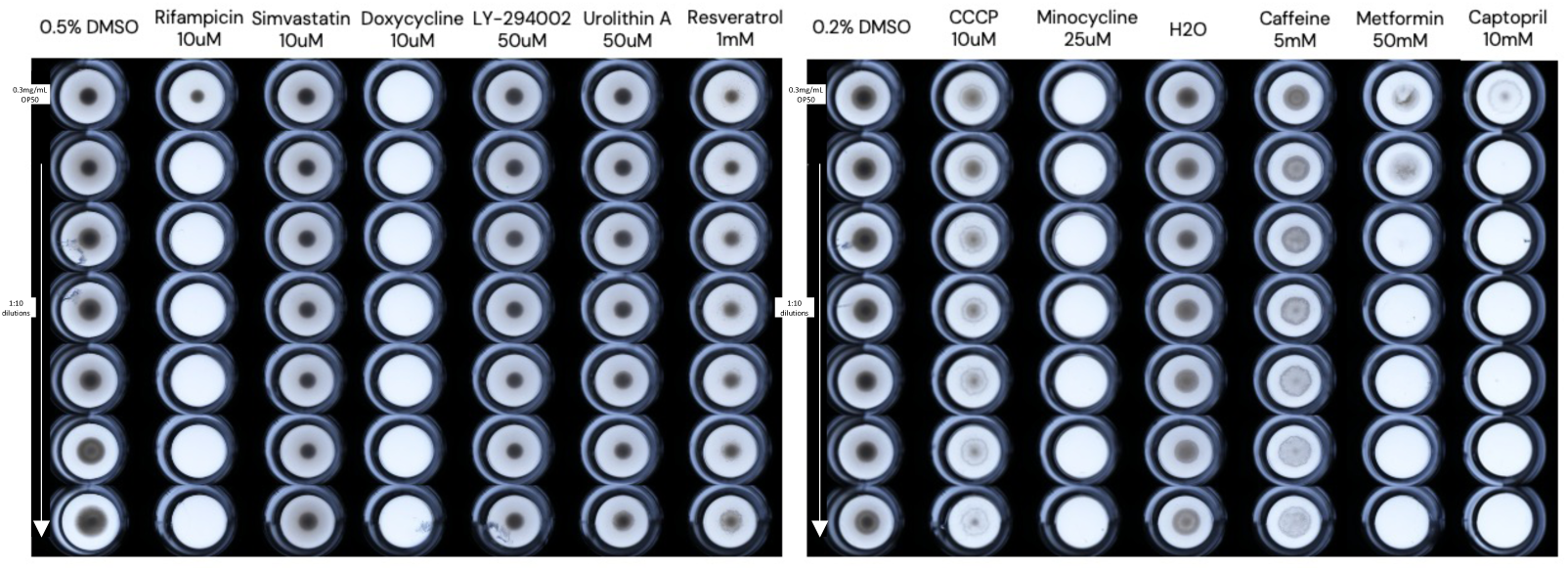
Direct effect of the drug on OP50 viability. Direct effect of the drug-concentration tested on the OP50 alone. Images of 7 1:10 serial dilutions starting from 0.3mg/mL OP50 after overnight exposure to Drug [Concentration] and pelleted.

**Figure S3.**
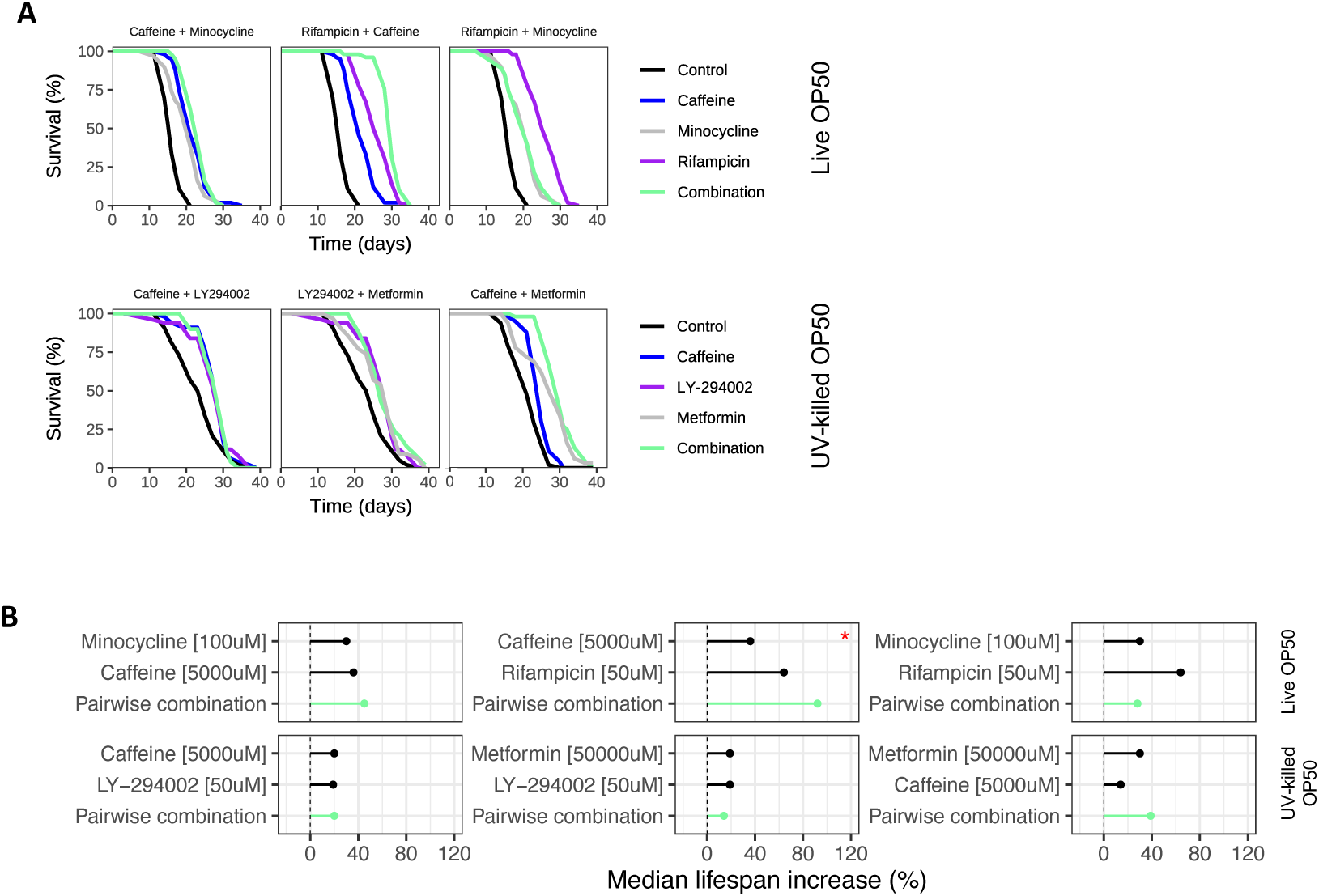
Combination of caffeine and rifampicin has a nearly additive impact on N2 lifespan when fed live OP50. (**A**) Survival curves for pairwise combinations of drugs tested on N2 lifespan when fed live OP50. Control in black, single drugs in blue, grey, purple, pairwise combination in light green. “*”: p < 0.05 for comparison of combination survival to all the single-drug results. (**B**) Summary effect of the single drugs and combinations on N2 lifespan from the survival curves. Median extension = 0% if insignificant (p ≥ 0.05). “*” indicates the combination result is significant compared to each single drug alone.

**Figure S4.**
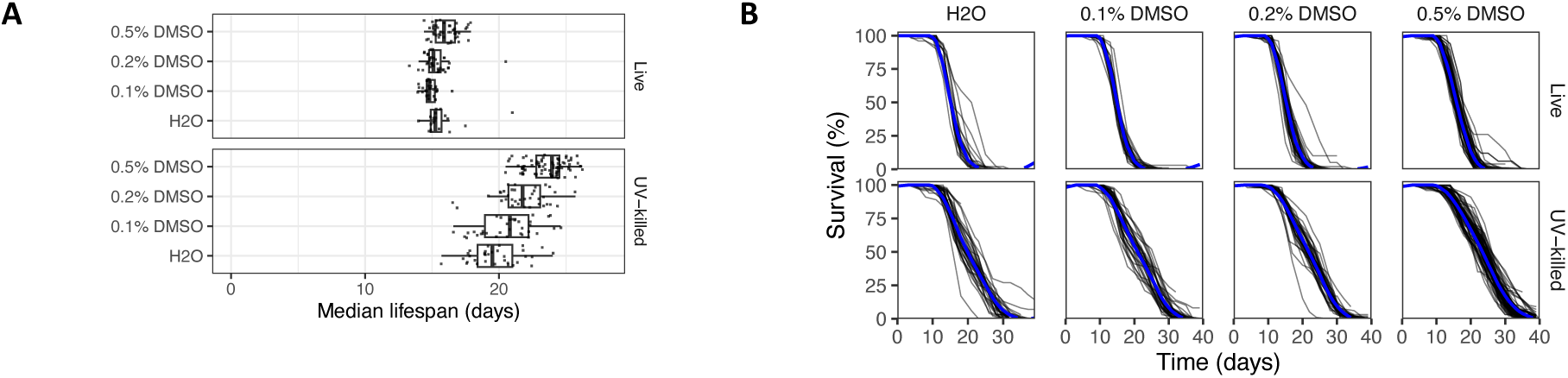
Variability is low among repeats of control experiments. (**A**) Boxplot of median survival from controls with 0%, 0.1%, 0.2%, or 0.5% DMSO vehicle concentrations, fed either live or UV-killed OP50. Each dot represents an individual control experiment. (**B**) Survival curves for each control experiment used in this study, fed either live or UV-killed OP50.

Supplemental Table 1. **Complete data for all lifespan experiments in this study.** Tabs with “Summary” contain the median result amongst trials (∼45 worms/trial) of the change in median survival at the best or all concentrations tested for each compound, separated by live or UV-killed OP50 diet. Median extension = 0 if insignificant (p ≥ 0.05). Tabs with “Figure” contain compound, concentration, solvent, N, and summary statistics for each individual trial for each figure in this study.

## References

1. Kenyon, C., Chang, J., Gensch, E., Rudner, A., & Tabtiang, R. (1993). A C. elegans mutant that lives twice as long as wild type. Nature, 366(6454), 461–464. 10.1038/366461a0

2. Friedman, D. B., & Johnson, T. E. (1988). A mutation in the age-1 gene in Caenorhabditis elegans lengthens life and reduces hermaphrodite fertility. Genetics, 118(1), 75–86. 10.1093/genetics/118.1.75

3. Wood, J. G., Rogina, B., Lavu, S., Howitz, K., Helfand, S. L., Tatar, M., & Sinclair, D. (2004). Sirtuin activators mimic caloric restriction and delay ageing in metazoans. Nature, 430(7000), 686–689. 10.1038/nature02789

4. Bass, T. M., Weinkove, D., Houthoofd, K., Gems, D., & Partridge, L. (2007). Effects of resveratrol on lifespan in Drosophila melanogaster and Caenorhabditis elegans. Mechanisms of Ageing and Development, 128(10), 546–552. 10.1016/j.mad.2007.07.007

5. Lucanic, M., Plummer, W. T., Chen, E., Harke, J., Foulger, A. C., Onken, B., Coleman-Hulbert, A. L., Dumas, K. J., Guo, S., Johnson, E., Bhaumik, D., Xue, J., Crist, A. B., Presley, M. P., Harinath, G., Sedore, C. A., Chamoli, M., Kamat, S., Chen, M. K., … Phillips, P. C. (2017). Impact of genetic background and experimental reproducibility on identifying chemical compounds with robust longevity effects. Nature Communications, 8(1), 14256. 10.1038/ncomms14256

6. Strong, R., Miller, R. A., Cheng, C. J., Nelson, J. F., Gelfond, J., Allani, S. K., Diaz, V., Dorigatti, A. O., Dorigatti, J., Fernandez, E., Galecki, A., Ginsburg, B., Hamilton, K. L., Javors, M. A., Kornfeld, K., Kaeberlein, M., Kumar, S., Lombard, D. B., Lopez-Cruzan, M., … Harrison, D. E. (2022). Lifespan benefits for the combination of rapamycin plus acarbose and for captopril in genetically heterogeneous mice. Aging Cell, 21(12), e13724. 10.1111/acel.13724

7. Lee, M. B., Blue, B., Muir, M., & Kaeberlein, M. (2023). The million-molecule challenge: a moonshot project to rapidly advance longevity intervention discovery. GeroScience, 1–11. 10.1007/s11357-023-00867-6

8. Barardo, D., Thornton, D., Thoppil, H., Walsh, M., Sharifi, S., Ferreira, S., Anžič, A., Fernandes, M., Monteiro, P., Grum, T., Cordeiro, R., De-Souza, E. A., Budovsky, A., Araujo, N., Gruber, J., Petrascheck, M., Fraifeld, V. E., Zhavoronkov, A., Moskalev, A., & Magalhães, J. P. de. (2017). The DrugAge database of aging-related drugs. Aging Cell, 16(3), 594–597. 10.1111/acel.12585

9. Wan, Q.-L., Zheng, S.-Q., Wu, G.-S., & Luo, H.-R. (2013). Aspirin extends the lifespan of Caenorhabditis elegans via AMPK and DAF-16/FOXO in dietary restriction pathway. Experimental Gerontology, 48(5), 499–506. 10.1016/j.exger.2013.02.020

10. Kumar, S., Dietrich, N., & Kornfeld, K. (2016). Angiotensin Converting Enzyme (ACE) Inhibitor Extends Caenorhabditis elegans Life Span. PLoS Genetics, 12(2), e1005866. 10.1371/journal.pgen.1005866

11. Lemire, B. D., Behrendt, M., DeCorby, A., & Gášková, D. (2009). C. elegans longevity pathways converge to decrease mitochondrial membrane potential. Mechanisms of Ageing and Development, 130(7), 461–465. 10.1016/j.mad.2009.05.001

12. Calvert, S., Tacutu, R., Sharifi, S., Teixeira, R., Ghosh, P., & Magalhães, J. P. de. (2016). A network pharmacology approach reveals new candidate caloric restriction mimetics in C. elegans. Aging Cell, 15(2), 256–266. 10.1111/acel.12432

13. Cabreiro, F., Au, C., Leung, K.-Y., Vergara-Irigaray, N., Cochemé, H. M., Noori, T., Weinkove, D., Schuster, E., Greene, N. D. E., & Gems, D. (2013). Metformin Retards Aging in C. elegans by Altering Microbial Folate and Methionine Metabolism. Cell, 153(1), 228–239. 10.1016/j.cell.2013.02.035

14. Bonuccelli, G., Brooks, D. R., Shepherd, S., Sotgia, F., & Lisanti, M. P. (2023). Antibiotics that target mitochondria extend lifespan in C. elegans. Aging (Albany NY), 15(21), 11764–11781. 10.18632/aging.205229

15. Solis, G. M., Kardakaris, R., Valentine, E. R., Bar-Peled, L., Chen, A. L., Blewett, M. M., McCormick, M. A., Williamson, J. R., Kennedy, B., Cravatt, B. F., & Petrascheck, M. (2018). Translation attenuation by minocycline enhances longevity and proteostasis in old post-stress-responsive organisms. ELife, 7, e40314. 10.7554/elife.40314

16. Golegaonkar, S., Tabrez, S. S., Pandit, A., Sethurathinam, S., Jagadeeshaprasad, M. G., Bansode, S., Sampathkumar, S., Kulkarni, M. J., & Mukhopadhyay, A. (2015). Rifampicin reduces advanced glycation end products and activates DAF-16 to increase lifespan in Caenorhabditis elegans. Aging Cell, 14(3), 463–473. 10.1111/acel.12327

17. Banse, S. A., Sedore, C. A., Johnson, E., Coleman-Hulbert, A. L., Onken, B., Hall, D., Jackson, E. G., Huynh, P., Foulger, A. C., Guo, S., Garrett, T., Xue, J., Inman, D., Morshead, M. L., Plummer, W. T., Chen, E., Bhaumik, D., Chen, M. K., Harinath, G., … Phillips, P. C. (2024). Antioxidants green tea extract and nordihydroguaiaretic acid confer species and strain-specific lifespan and health effects in Caenorhabditis nematodes. GeroScience, 46(2), 2239– 2251. 10.1007/s11357-023-00978-0

18. Robida-Stubbs, S., Glover-Cutter, K., Lamming, D. W., Mizunuma, M., Narasimhan, S. D., Neumann-Haefelin, E., Sabatini, D. M., & Blackwell, T. K. (2012). TOR Signaling and Rapamycin Influence Longevity by Regulating SKN-1/Nrf and DAF-16/FoxO. Cell Metabolism, 15(5), 713–724. 10.1016/j.cmet.2012.04.007

19. Jahn, A., Scherer, B., Fritz, G., & Honnen, S. (2020). Statins Induce a DAF-16/Foxo-dependent Longevity Phenotype via JNK-1 through Mevalonate Depletion in C. elegans. Aging and Disease, 11(1), 60–72. 10.14336/ad.2019.0416

20. Liu, Y. J., Janssens, G. E., McIntyre, R. L., Molenaars, M., Kamble, R., Gao, A. W., Jongejan, A., Weeghel, M. van, MacInnes, A. W., & Houtkooper, R. H. (2019). Glycine promotes longevity in Caenorhabditis elegans in a methionine cycle-dependent fashion. PLoS Genetics, 15(3), e1007633. 10.1371/journal.pgen.1007633

21. Sutphin, G. L., Bishop, E., Yanos, M. E., Moller, R. M., & Kaeberlein, M. (2012). Caffeine extends life span, improves healthspan, and delays age-associated pathology in Caenorhabditis elegans. Longevity & Healthspan, 1(1), 9–9. 10.1186/2046-2395-1-9

22. Chen, W., Rezaizadehnajafi, L., & Wink, M. (2013). Influence of resveratrol on oxidative stress resistance and life span in Caenorhabditis elegans. Journal of Pharmacy and Pharmacology, 65(5), 682–688. 10.1111/jphp.12023

23. Ryu, D., Mouchiroud, L., Andreux, P. A., Katsyuba, E., Moullan, N., Nicolet-dit-Félix, A. A., Williams, E. G., Jha, P., Sasso, G. L., Huzard, D., Aebischer, P., Sandi, C., Rinsch, C., & Auwerx, J. (2016). Urolithin A induces mitophagy and prolongs lifespan in C. elegans and increases muscle function in rodents. Nature Medicine, 22(8), 879–888. 10.1038/nm.4132

24. Zhao, Y., Gilliat, A. F., Ziehm, M., Turmaine, M., Wang, H., Ezcurra, M., Yang, C., Phillips, G., McBay, D., Zhang, W. B., Partridge, L., Pincus, Z., & Gems, D. (2017). Two forms of death in ageing Caenorhabditis elegans. Nature Communications, 8(1), 15458. 10.1038/ncomms15458

25. Gems, D., & Riddle, D. L. (2000). Genetic, Behavioral and Environmental Determinants of Male Longevity in Caenorhabditis elegans. Genetics, 154(4), 1597–1610. 10.1093/genetics/154.4.1597

26. Bennett, D. F., Goyala, A., Statzer, C., Beckett, C. W., Tyshkovskiy, A., Gladyshev, V. N., Ewald, C. Y., & Magalhães, J. P. de. (2023). Rilmenidine extends lifespan and healthspan in Caenorhabditis elegans via a nischarin I1-imidazoline receptor. Aging Cell, 22(2), e13774. 10.1111/acel.13774

27. Aiello, G., Sabino, C., Pernici, D., Audano, M., Antonica, F., Gianesello, M., Ballabio, C., Quattrone, A., Mitro, N., Romanel, A., Soldano, A., & Tiberi, L. (2022). Transient rapamycin treatment during developmental stage extends lifespan in Mus musculus and Drosophila melanogaster. EMBO Reports, 23(9), e55299. 10.15252/embr.202255299

28. Janssens, G. E., Lin, X.-X., Millan-Ariño, L., Kavšek, A., Sen, I., Seinstra, R. I., Stroustrup, N., Nollen, E. A. A., & Riedel, C. G. (2019). Transcriptomics-Based Screening Identifies Pharmacological Inhibition of Hsp90 as a Means to Defer Aging. Cell Reports, 27(2), 467–480.e6. 10.1016/j.celrep.2019.03.044

29. Ye, X., Linton, J. M., Schork, N. J., Buck, L. B., & Petrascheck, M. (2014). A pharmacological network for lifespan extension in Caenorhabditis elegans. Aging Cell, 13(2), 206–215. 10.1111/acel.12163

30. Zhou, C., Zhou, Y., Liang, Y., Chen, L., Liu, L., Wei, F., & Li, G. (2023). Fluoxetine Promotes Longevity via Reactive Oxygen Species in Caenorhabditis elegans. The Journals of Gerontology: Series A, 79(1), glad220. 10.1093/gerona/glad220

31. Spindler, S. R., Li, R., Dhahbi, J. M., Yamakawa, A., & Sauer, F. (2012). Novel Protein Kinase Signaling Systems Regulating Lifespan Identified by Small Molecule Library Screening Using Drosophila. PLoS ONE, 7(2), e29782. 10.1371/journal.pone.0029782

32. Lithgow, G. J., Driscoll, M., & Phillips, P. (2017). A long journey to reproducible results. Nature, 548(7668), 387–388. 10.1038/548387a

33. Urban, N. D., Cavataio, J. P., Berry, Y., Vang, B., Maddali, A., Sukpraphrute, R. J., Schnell, S., & Truttmann, M. C. (2021). Explaining inter-lab variance in C. elegans N2 lifespan: Making a case for standardized reporting to enhance reproducibility. Experimental Gerontology, 156, 111622. 10.1016/j.exger.2021.111622

34. Harrison, D. E., Strong, R., Sharp, Z. D., Nelson, J. F., Astle, C. M., Flurkey, K., Nadon, N. L., Wilkinson, J. E., Frenkel, K., Carter, C. S., Pahor, M., Javors, M. A., Fernandez, E., & Miller, R. A. (2009). Rapamycin fed late in life extends lifespan in genetically heterogeneous mice. Nature, 460(7253), 392–395. 10.1038/nature08221

